# Gene discovery and functional assessment of rare copy-number variants in neurodevelopmental disorders

**DOI:** 10.1101/011510

**Authors:** Janani Iyer, Santhosh Girirajan

## Abstract

Rare copy-number variants (CNVs) are a significant cause of neurodevelopmental disorders. The sequence architecture of the human genome predisposes certain individuals to deletions and duplications within specific genomic regions. While assessment of individuals with different breakpoints has identified causal genes for certain rare CNVs, deriving gene-phenotype correlations for rare CNVs with similar breakpoints has been challenging. We present a comprehensive review of the literature related to genetic architecture that is predisposed to recurrent rearrangements, and functional evaluation of deletions, duplications, and candidate genes within rare CNV intervals using mouse, zebrafish, and fruit fly models. It is clear that phenotypic assessment and complete genetic evaluation of large cohorts of individuals carrying specific CNVs and functional evaluation using multiple animal models are necessary to understand the molecular genetic basis of neurodevelopmental disorders.

## KEY POINTS

- Genetic architecture predisposes about 9% of the human genome to disease-causing rare deletions and duplications
- Chromosomal translocations and atypical deletions have enabled the discovery of dosage sensitive genes within rare CNV regions
- Mouse models carrying deletions and duplications for rare CNVs have recapitulated most of the human disease phenotypes and have also aided gene discovery in disorders where there are limitations to human studies
- Zebrafish studies have proven to be excellent tools for high throughput screens and for evaluation of key neuroanatomical phenotypes
- *Drosophila* models have provided deeper insights into the neuronal basis of neurodevelopmental disorders
- Future studies of rare CNVs should consider the limitations of animal models while emphasizing validation across model systems for accurate understanding of the pathophysiology of human disease

## INTRODUCTION

Copy-number variants (CNVs) are a significant source of genetic variation and have an important role in human variation, disease, and evolution [1]. In terms of the number of base pairs, between any two individuals, there is more variation due to CNVs, including deletions and duplications than SNPs [2]. Rare CNVs typically occur at a frequency of <1% to <0.1% in the population [3]. Recent high throughput studies using microarrays and sequencing have provided insights into the structure of the human genome [4]. The following themes have emerged from these studies. First, sequence architecture of the human genome predisposes certain regions to recurrent and non-recurrent rearrangements. Genetic predisposition for recombination-based events is conferred by a specific configuration of repeat sequences facilitated by large inversions. In addition, smaller repeats have also led to non-recurrent rearrangements through errors in recombination and replication. Second, rare CNVs are particularly enriched in individuals with neuropsychiatric and developmental phenotypes and overlapping CNVs with different breakpoints have narrowed down the candidate genes for specific phenotypes. Third, while animal models have been able to recapitulate dosage-sensitive phenotypes for one or more genes within certain CNVs, for others, prioritization of candidate genes has been difficult because of a lack of gene-specific chromosomal translocations or atypical events narrowing down the CNV critical interval to individual genes.

## Genetic architecture predisposes individuals to recurrent and non-recurrent CNVs

Local sequence architecture predisposes certain regions of the genome to recurrent rearrangements [5–7]. While an estimated 69% of these events occur as a result of recombination errors during meiosis, other mechanisms including non-homologous end joining, replication-based events (such as microhomology-mediated break-induced replication or MMBIR/fork stalling and template switching or FoSTeS), and complex rearrangements due to multiple template switching events have also been reported [8]. Non-allelic homologous recombination (NAHR), classically termed unequal crossover was originally proposed for copy number changes at the *Bar* locus in fruit flies [9, 10], and was also used to explain the mechanism for gene loss and gain at the hemoglobin alpha subunit (HBA) [11], and evolution and expansion of repeat sequences in yeast [12] and humans [13].

For disorders resulting from genomic rearrangements, the concept of NAHR was initially described where deletions and duplications of the genomic interval on chromosome 17p12 containing *PMP22* were associated with Charcot-Marie-Tooth disease 1A (CMT1A) and Hereditary neuropathy with liability to pressure palsies (HNPP), respectively [14, 15]. These genomic intervals are flanked by large blocks of repeats with high sequence identity called segmental duplications (SDs) or low copy repeats that form substrates for NAHR events [5, 16]. The products of these events will either be a deletion or a reciprocal duplication leading to haploinsufficiency or triploinsufficiency of genes within the region. This signature feature predisposes up to 9% of the human genome to recurrent deletions and duplications making approximately 3000 genes vulnerable to dosage alteration [17]. A genome-wide assessment revealed as many as 130 such hotspots, which are flanked by SDs in the human genome that predisposes individuals to neurodevelopmental disorders [7, 17]. Other classical examples of disorders caused by SD-mediated rearrangements include Smith-Magenis syndrome (SMS) [18] and Potocki-Lupski syndrome (PTLS) [19] (17p11.2 deletion and duplication respectively), Williams-Beuren (WBS) syndrome (7q11.23 deletion) [20] and 7q11.23 duplication [21], Sotos syndrome (5q35 deletion) [22], DiGeorge/velocardiofacial syndrome (DGS) (22q11.2 deletion) [23], Prader-Willi (PWS)/Angelman syndrome (AS) (15q11.2-q13.1 deletion) [24], and 17q21.1 deletion [7]. While most of the classically defined syndromes were identified through conventional cytogenetics [25] and careful phenotypic assessment, recent studies using targeted approaches for novel CNV discoveries have led to the identification of another class of CNVs that are more common in frequency and are associated with a range of neurodevelopmental phenotypes including autism, intellectual disability, epilepsy, and schizophrenia. Examples in this series include CNVs within 1q21.1, 3q29, 15q13.3, 15q24, 16p11.2, 16p12.2, 16p13.11, and 17q12 [26].

Genetic predilections to rearrangements within hotspots are dependent upon the presence of directly oriented SDs of high sequence identity [5]. Such configurations are present at different frequencies in the population making only a subset of the individuals susceptible, while others without the predisposing architecture are protected against disease-associated rearrangements [27, 28]. The density of SDs seems to correlate with susceptibility to rearrangements [29]. For example, increased copy numbers of SDs flanking the 7q11.23 region have been linked to a pronounced risk for rearrangements leading to WBS [30]. Structural susceptibility to rearrangements is particularly favored by inversions in the region, which align the two participating SDs in direct orientation. Inversion polymorphisms have been described for several human disease-associated regions including chromosomal regions 3q29 [31], 7q11.23 [32], 8p23 [33], 15q11.2q13.1 [34], 15q13.3 [35], 15q24.1, 16p12.1, 17q12, and 17q21.31 [36]. For example, SD architecture at the recently described 16p12.1 microdeletion has two structural configurations, referred to as the S1 and S2 haplotypes [27]. The two haplotypes have different worldwide frequencies (frequency for S1 is 17.6% and S2 is 82.4%) and differ by 333 kbp of additional sequence in S2 compared to S1. In the rearrangement prone S2 configuration there is a direct orientation of specific SDs flanking the disease-associated region, which is not seen in the protective S1 haplotype. Similar analyses have been performed for other hotspot regions providing a map of regions that are susceptible or protective against recurrent rearrangements [31, 37].

Replication-based mechanisms such as FoSTeS or MMBIR have been proposed to explain non-recurrent and complex genomic rearrangements [38, 39]. These models were described in human disease-associated rearrangements based on the observations in yeast [40] and bacteria [41] that have shown similar mechanistic signatures [42]. These rearrangements are generally characterized by multiple rounds of template switching over long distances leading to multiple copy number changes and the presence of microhomology at breakpoint junctions. The FoSTeS/MMBIR mechanism has been implicated in *PLP1* duplications in Pelizaeus-Merzbacher disease [38], *MECP2* duplication [43], *FOXL2* deletions [44], and *LIS1* deletions and duplications [42]. From these studies it is clear that multiple mechanisms based on DNA recombination, replication, and repair processes are involved in genome evolution.

## Causal gene discovery within rare CNVs

Rare CNVs often contain multiple unrelated genes and manifest complex neurodevelopmental and neuropsychiatric phenotypes [45]. Due to the large number of genes involved, it has been challenging to identify specific genes of functional relevance and correlate them to a phenotype. Gene discovery within rare CNVs has been mostly aided by chromosomal events unraveling the causative genes involved in a specific phenotype. Serendipitous translocations disrupting causal genes within the region have facilitated the discovery of *NSD1* in Sotos syndrome [46], and *SHANK3* in Phelan-McDermid syndrome [47]. Similarly, a maternally inherited inversion led to the discovery of *UBE3A* in Angelman syndrome [48]. Mapping of CNVs from patients displaying the same phenotypes but with atypical deletions or duplications has been used in identifying the minimum critical region carrying the candidate genes. Examples include the narrowing of the 17q21.31 region to a single gene by using atypical events that were not mediated by NAHR, leading to the identification of *KANSL1* (involved in histone acetylation) as a causative gene in 17q21.31 microdeletion syndrome [49, 50]. In fact, clinical features in individuals with *KANSL1* mutations were almost indistinguishable from those carrying the deletion. Another example is the use of atypical deletions caused by alternate pairs of segmental duplications to identify a critical region containing three genes, *RAI1*, *TOM1L2*, and *DRG2*, within the SMS deletion interval [51, 52]. Further sequencing of these genes in individuals manifesting most SMS features resulted in the identification of disruptive mutations in the retinoic acid induced 1 (*RAI1*) gene. Atypical duplications within 17p11.2 also pointed towards *RAI1* as a major gene for PTLS [53]. This approach is limited by the availability of patients with similar phenotypes carrying atypical deletions and is more geared towards identifying loss of function alleles than gain of function. In a recent study, Coe and colleagues extended the morbidity map of CNVs in the human genome by analyzing 29,085 children with developmental delay and 19,584 healthy controls. Comparing enrichment of deletions and duplications in the two groups over the entire genome, this study identified 70 regions significantly associated with disease [54]. They were also able to identify several novel and atypical CNVs pointing to specific genes, including 26 candidates that were further interrogated for loss of function mutations within 4,716 cases and 2,193 controls by targeted sequencing approaches. By integrating the analysis of CNVs and single nucleotide mutations, ten genes associated with specific subtypes of developmental delay disorders were identified.

Unmasking of recessive alleles has also contributed to identifying the causal genes for specific phenotypes. Recessive alleles manifest their phenotype when they are in a homozygous state. However, a recessive allele on one chromosome can be unmasked to display its phenotype by deletion of the normal allele on the other chromosome. Thus the patient will exhibit phenotypes resulting from the haploinsufficiency of dosage-sensitive genes in the deleted region and from loss-of-function due to the recessive mutation. For example, a recessive mutation in the myosin gene (*MYO15A*) within the 17p11.2 region was identified in a SMS patient manifesting profound sensorineural hearing loss [55]. In a recent study of an individual with the autism/developmental delay associated 16p11.2 deletion, the clinical phenotypes were different from the usual 16p11.2 deletion phenotypes, and chromosomal microarray analysis revealed an additional 220-kbp deletion of the distal 16p11.2 region [56]. Since the affected individual’s phenotypes were similar to juvenile neuronal ceroid-lipofuscinosis or Batten disease and the 220-kbp deletion partially included *CLN3*, a known candidate gene for the phenotype, Pebrel-Richard and colleagues were able to sequence and confirm a mutation in the *CLN3* allele. Unmasking of the recessive *CLN3* mutation therefore explained the variability observed in this patient. Similarly, oculocutaneous albinism was reported in an individual with Prader-Willi syndrome (PWS) and a recessive mutation of *OCA2*, a gene essential for normal pigmentation [57]. A complex inheritance pattern was described for individuals carrying proximal deletions of 1q21.1 and manifesting distal thrombocytopenia-absent radius (TAR) syndrome. In contrast to other syndromic disorders, where almost all CNVs are *de novo*, this deletion was also observed in unaffected parents suggesting the deletion to be necessary but not sufficient to cause the syndrome [58]. Further studies using exome sequencing of TAR patients with 1q21.1 deletions identified low-frequency non-coding SNPs located in the 5’ UTR and first intron of *RBM8A* and follow up *in vitro* studies revealed that these non-coding SNPs caused a decrease in *RBM8A* expression [59]. These studies suggested a compound or biallelic inheritance in TAR syndrome of a non-coding SNP and a rare null allele in *RBM8A* (1q21.1 microdeletion).

## Models to explain causality of disease due to rare CNVs

The association of genes in rare CNVs with phenotypes can be broadly categorized under three models (Figure 1). In the single gene model there is one major gene that leads to most of the disease phenotypes. An example from this category is the retinoic acid induced 1 (*RAI1*) gene implicated in Smith-Magenis syndrome. Comparison of clinical features of children with the deletion with those with *RAI1* mutations suggested that haploinsuficiency of *RAI1* is responsible for the core features of SMS while the variability and severity of the disorder was conferred by other genes in the deletion region [60]. Similarly, the cardinal features of Sotos syndrome were observed in similar proportions in individuals with *NSD1* mutations compared to those with 5q35 deletions [61]. Thus, the major features of Sotos syndrome can be explained by haploinsufficiency of *NSD1*. Evidence from translocation breakpoints and atypical deletions pointed to *EHMT1* as a major contributor towards the clinical features observed in individuals with 9q34 deletion or Kleefstra syndrome [62]. Further, patients without the 9q34 deletion but with loss-of-function *EHMT1* mutations showed typical features of Kleefstra syndrome, including severe intellectual disability, hypotonia, pathognomonic craniofacial features, conotruncal heart defects, and behavioral problems, exemplifying the one gene model for rare CNV disorders [63].

**Figure 1.**
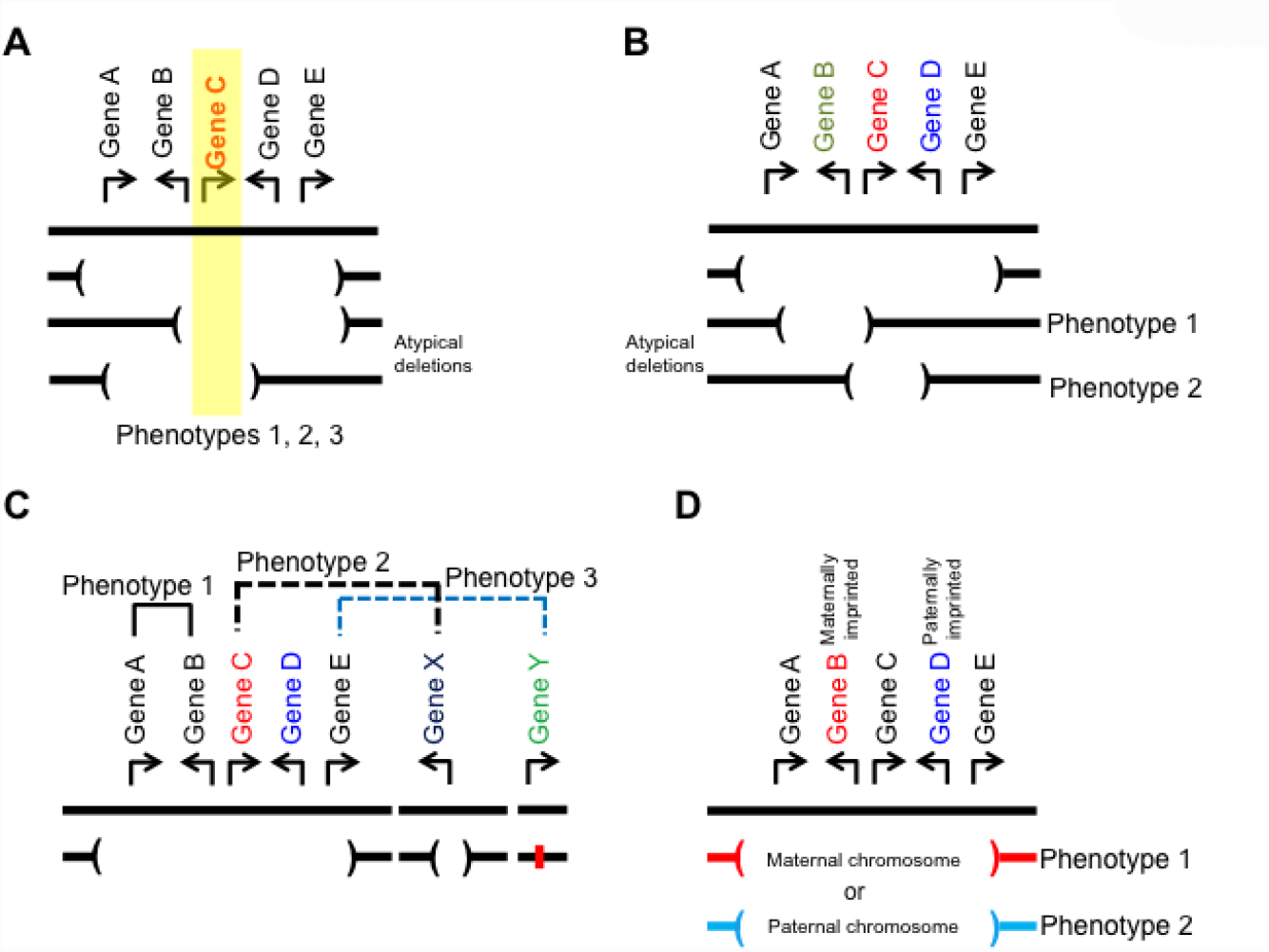

Models to explain causality of disease due to rare CNVs. **(A)** Single gene model: A core gene is responsible for a majority of the disease phenotypes as observed in SMS. **(B)** Contiguous gene model: Individual genes within the CNV region contribute to a specific phenotype and as a result individuals with the deletion manifest multiple unrelated features as in WBS. **(C)** *Cis* and *trans* interaction model: Dosage alteration of two or more functionally relevant genes within (*cis*) or outside the CNV region (*trans*) leads to the disease phenotype. **(D)** Parent-of-origin model: Disease phenotypes due to disruption of the paternally or the maternally derived allele/genes as in PWS and AS, respectively.

In the contiguous gene model, each gene contributes to a phenotype, which collectively results in multiple unrelated features in a single individual carrying the rare CNV. A classical example is that of WBS, which is associated with a constellation of clinical features including facial dysmorphism, intellectual disability, connective tissue abnormalities, supravalvular aortic stenosis, hypersociability, hypercalcemia, and behavioral features [64]. Chromosomal deletions on 7q11.23 leading to WBS results in the disruption of 26 to 28 genes. Contribution of haploinsufficiency of certain individual genes towards specific phenotypes in WBS was essentially identified by analysis of atypical deletions. Typical characteristic features of WBS include facial dysmorphism, connective tissue alterations, mild growth retardation, cardiac defects, mild-to-moderate intellectual disability and behavioral defects including motor problems, hypersociability and visuospatial defects [64]. Multiple studies involving meticulous phenotyping of each of the WBS phenotypes in combination with mapping of atypical deletions resulted in the correlation of specific phenotypes to genes within 7q11.23. For example, haploinsufficiency of *ELN* was implicated in cardiac phenotypes [65], *GTF2IRD1* for craniofacial features, *GTF2I* and *GTF2IRD1* for neurobehavioral features [66–68], and *LIMK1* for visuospatial impairments. Interestingly, haploinsufficiency of *NCF1* was identified to be protective against hypertension in individuals with the deletion [69]. Further analysis of atypical deletions extending telomeric to the WBS region by Marshall and colleagues implicated *MAGI2* as causal for infantile spasms [70].

In the *cis*-interaction model, the combined effect of dosage alteration of two or more genes within the CNV region leads to the disease phenotypes. Theoretically, phenotypes caused by CNVs can be expected to be due to additive or synergistic interaction of multiple genes within the region. This model can potentially explain the severity of brain malformations in individuals carrying 17p13.3 deletions containing both *PAFAH1B1* (encoding LIS1) and *YWHAE* [71]. Individuals with deletions involving *PAFAH1B1* present with isolated lissencephaly features while individuals with larger deletions involving *YWHAE* show more severe forms of lissencephaly. Using mouse studies, Toyo-oka and colleagues showed that *YWHAE* is an essential component for optimal neuronal migration and acted through its interaction with NUDEL, a protein that also interacts with LIS1 for neuronal migration [72]. Thus, disruption of two genes interacting within the same multi-protein complex required for neuronal migration resulted in a more severe phenotype. A recent example of the *cis*-interaction model was observed for 17p13.1 deletion syndrome where dosage alterations of discrete pairings of genes within the region induced microcephaly phenotypes [73].

The concept of genetic background and modifier genes has been proposed previously [74, 75]. It was recently observed that the phenotypic effect of rare CNVs could be modified by another large CNV elsewhere in the genome. The *trans*-interaction model can be illustrated by the 16p12.1 microdeletion. Severely affected children often inherited the deletion from a carrier parent who manifested only subtle neuropsychiatric features [76]. Further investigation revealed that about 25% of the probands carried another large CNV elsewhere in the genome, in addition to the 16p12.1 deletion, potentially dictating the ultimate phenotypic outcome. Extension of this “two-hit” model suggested a generalizable feature for rare CNVs [77]. Individuals manifesting CNV syndromes such as Sotos syndrome and Smith-Magenis syndrome carried fewer or no second hits while those carrying CNVs associated with variable expressivity such as 16p11.2 deletion, 15q13.3 deletion, and 1q21.1 deletion carried another large CNV. The *trans*-interaction model could explain these observations, wherein there is a compounding effect of two or more genes interacting in the same or related pathways in an additive or synergistic manner [78].

In the parent-of-origin model, phenotypes can vary depending upon whether the disrupted genes were derived from maternal or paternal chromosomes. Loss of the paternal segment of chromosomal region 15q11.2-q13.1 results in PWS, characterized by growth retardation, hypotonia, intellectual disability, obesity, hypogonadism, craniofacial features, and short stature [79]. Originally described as a true contiguous gene syndrome, recent studies have identified patients with loss of only one of the snoRNA clusters within the paternally imprinted segment who still manifest typical features of PWS [80, 81]. Similarly, loss of the maternal segment of ubiquitin-protein ligase E3A (*UBE3A*) within 15q11.2-q13.1 results in AS with individuals manifesting intellectual disability, speech and language delay, movement disorders, behavioral abnormalities, and seizures [48, 82].

## Animal models for rare CNV disorders

Animal models for human diseases are important for validation of pathogenicity, establishing a link between genes/genomic regions and phenotypes, and providing a resource to study disease mechanisms relevant to human physiology. Animal models have been especially useful in delineating specific genes within disease-associated regions that were not mapped by translocations or atypical CNVs. We will focus on examples from mouse, zebrafish, and fruit fly models utilized for functional evaluation of rare CNV disorders (Table 1).

**Table 1:** A summary of clinical features and functional studies of rare CNV disorders.

| Genomic disorder | Clinical features | Candidate gene(s) | Animal Model | Phenotypes in animal models | Reference |
| --- | --- | --- | --- | --- | --- |
| Williams-Beuren Syndrome (7q11.23 deletion); OMIM: 194050 | Characteristic facial features, cardiac malformation, cognitive deficits, behavioral defects, hypersociability | Contiguous gene syndrome | Mouse (complete Deletion) | Mice exhibit reduced body weight, craniofacial and brain abnormalities, increased sociability, cognitive deficits, mild cardiovascular phenotype, reduced brain size and motor defects | [117] |
|  |  |  | Mouse (proximal deletion, lacking *Gtf2i* to *Limk1*) | Increased sociability and acoustic startle response and retarded growth | [112] |
|  |  |  | Mouse (distal deletion, lacking *Limk1* to *Fkbp6*) | Cognitive deficits, shortened skull and reduced brain size | [112] |
|  |  |  | Mouse (proximal plus distal to achieve complete deletion) | Growth retardation, shortened skull, smaller brains and impaired motor skills | [112] |
|  |  |  | Mouse (*Wstf*<sup>-/-</sup>) | Neonatal lethality and cardiovascular abnormalities | [113] |
|  |  |  | Mouse (*Eif4h* knockout) | Growth retardation, reduced brain size, altered brain morphology, impaired memory and behavioral defects | [114] |
|  |  |  | Mouse (partial deletion lacking *Cyln2*) | Mild growth defects, brain abnormalities, hippocampal dysfunction and motor defects | [115] |
|  |  |  | Mouse (*Eln* knockout) | Die of obstructive arterial disease mimicking atherosclerosis | [116] |
|  |  |  | Mouse (*Gtf2i*<sup>-/-</sup>, *Gtf2i*<sup>+/-</sup>) | Craniofacial and neurobehavioral abnormalities. *Gtf2i*<sup>-/-</sup> mice show embryonic lethality and infertility | [136] |
|  |  |  | Mouse (*Gtf2ird1*<sup>-/-</sup>) | Craniofacial abnormalities | [68] |
| 7q11.23 duplication; OMIM: 609757 | Mild craniofacial anomalies, speech delay, congenital abnormalities, ID, autism and schizophrenia | Contiguous gene syndrome | Mouse (complete duplication) | Not available yet |  |
|  |  |  | Mouse (*Gtf2i* duplication) | Increased maternal-separation anxiety | [118] |
| Smith-Magenis Syndrome (17p11.2 deletion); OMIM: 182290 | ID, dysmorphic features, congenital anomalies, sleep disturbance, self-injurious and aggressive behavior | *RAI1* | Mouse (complete deletion) | Mice exhibit obesity, craniofacial abnormalities, seizures and abnormal behavior | [105] |
|  |  |  | Mouse (*Rai1* +/-, *Rai1* -/-) | *Rai1* +/- mice display obesity and craniofacial features. *Rai1* -/- show embryonic lethality and postnatal growth retardation | [104] |
| Potocki-Lupski Syndrome (17p11.2 duplication); OMIM: 610883 | DD, hypotonia, failure to thrive, ID, cardiovascular malformations, behavioral abnormalities including ASD | *RAI1* | Mouse (complete duplication) | Reduced weight, hyperactivity, abnormal social behaviors | [105, 111] |
|  |  |  | Mouse (overexpression of *Rai1*) | Growth retardation, increased locomotor activity, and abnormal anxiety-related behavior | [109, 110] |
| Angelman Syndrome (15q11-q13 maternal deletion); OMIM: 105830 | Intellectual disability, ataxia, developmental delay, behavioral defects and speech and language impairments | *UBE3A* | Mouse (*Ube3A* knockout) | *Ube3A* maternal-deficient mice display reduced brain weight, ataxia, motor defects and behavioral defects | [137, 138] |
|  |  |  | *Drosophila* | Lack of *Ube3A* in *Drosophila* leads to motor defects, abnormal circadian rhythms, long-term memory defects and abnormalities in dendritic arborization in the peripheral nervous system | [133, 134] |
| Prader-Willi Syndrome (15q11-q13 paternal deletion); OMIM: 176270 | Facial dysmorphism, hypogonadism, short stature, obesity, ID, and behavioral problems | Contiguous gene syndrome | Mouse (*TgPWS*: transgene inserted into chromosome 7C) | Paternal transmission results in a lack of paternal gene expression of the PWS interval that leads to early postnatal lethality | [139] |
|  |  |  | Mouse (PWS-IC +/- lacking the 42 kb PWS imprinting center including *Snrpn*, *Znf127*, *Ndn*, *Ipw*) | Failure to thrive and early postnatal lethality | [140] |
|  |  |  | Mouse (PWS-IC lacking a smaller 6 kb PWS imprinting center including *Snrpn*) | Mice with paternal inheritance of this deletion exhibit complete loss of paternal gene expression and neonatal lethality | [141] |
|  |  |  | Mouse (*Magel2* knockout) | Failure to survive to adulthood | [142] |
|  |  |  | Mouse (*Snord116* knockout) | Early-onset postnatal growth deficiency, normal fertility, increased anxiety, defects in motor learning | [143] |
|  |  |  | Mouse (*Ndn* knockout) | *Ndn* knockout mice survive and display increased skin scraping and improved spatial learning and memory | [144] |
| 15q11-q13 duplication; OMIM: 608636 | Autism, intellectual disability, ataxia, seizures, developmental delays, and behavioral problems |  | Mouse (complete duplication) | The mice with paternally inherited duplication exhibited autistic behaviors such as poor social interaction and abnormal ultrasonic vocalizations, whereas the maternally inherited duplication mice were WT-like | [145] |
|  |  |  | Mouse (*Ube3A* overexpression) | Autism like phenotypes and decreased glutamate synaptic transmission, irrespective of the paternal origin. However the *Ube3A* transgene was not imprinted | [146] |
| 16p11.2 deletion; OMIM: 611913 | ID, ASD, obesity, DD, speech delays |  | Mouse (complete deletion) | 50% mice die postnatally and the survivors display abnormalities in brain architecture, increased brain volume and behavioral defects | [119] |
|  |  |  | Mouse (complete deletion) | Defects in basal ganglia circuitry, synaptic defects, elevated numbers of medium spiny neurons and behavioral defects including hyperactivity and cognitive deficits | [120] |
|  |  |  | Zebrafish | Suppression of *KCTD13* mimics the macrocephaly phenotype seen in patients. | [125] |
| 16p11.2 duplication; OMIM: 614671 | ID, ASD, schizophrenia |  | Mouse (complete duplication) | 16p11.2 duplication mice display milder phenotypes as compared to the deletion mice. These mice exhibit reduced brain volume and mild behavioral defects. | [119] |
|  |  |  | Zebrafish | Overexpression of *KCTD13* mimics the microcephaly phenotype seen in patients. | [125] |
| DGS/VCFS (22q11.2 deletion); OMIM: 188400; OMIM: 192430 | Craniofacial abnormalities, cardiovascular defects, ID, ASD, and schizophrenia | *TBX1* | Mouse (Df/+ carrying deletion of 22 genes within 22q11.2) | Df1/+ mice exhibit cardiovascular defects, learning and memory deficits and abnormal behavior, as measured by prepulse inhibition of the startle response | [91, 94] |
|  |  |  | Mouse (overlapping deletions within the 22q11.2 region and *Tbx1* knockouts) | Overlapping deletions in mouse models narrowed down the intervals responsible for PPI impairments to *Tbx1* and *Gnb1l* and for heart defect to *Tbx1*. These were further validated by targeted knock-out of *Tbx1*. | [88, 89, 91, 94] |
| Phelan-McDermid syndrome; (22q13 deletion); OMIM: 606232 | ID, DD, delayed or absent speech, and autistic features | *SHANK3* | Mouse (*Shank3*+/-) | Defects in synaptic function and plasticity and reduced social interactions. | [147] |
|  |  |  | Mouse (*Shank3*-/-) | Abnormal social behavior, repetitive behavior, defects in learning and memory, morphological alterations in dendritic spines, and impaired long term potentiation | [148, 149] |
| Brachydactyly Mental Retardation Syndrome (2q37 deletion); OMIM: 600430 | ID, DD, obesity, craniofacial and skeletal defects, behavioral defects, sleep disturbance and ASD | *HDAC4* | Mouse (*Hdac4*-/-) | Severe bone malformations as indicated by chondrocyte hypertrophy and premature bone ossification | [150] |
| Kleefstra Syndrome; (9q34 del); OMIM: 610253 | Severe intellectual disability, hypotonia, brachycephaly, craniofacial abnormalities, conotruncal heart defects, and behavioral problems | *EHMT1* | Mouse (*Ehmt*+/-) | Autistic like features, delayed postnatal development, hypotonia, craniofacial dysmorphism and defects in learning and synaptic dysfunction | [151] |
|  |  |  | *Drosophila* | *EHMT* deficient flies show abnormal dendritic arbors, defects in larval locomotor and non-associative learning and courtship memory | [135] |
| 15q13.3 deletion;<br>OMIM: 612001 | Idiopathic generalized epilepsy, ID, ASD and schizophrenia | *OTUD7*,<br>*CHRNA7* | Mouse (complete deletion) | Susceptibility to seizures, impaired spatial reference memory, aggression, increased body weight and auditory processing defects | [121] |
|  |  |  | Mouse<br>(*Chrna7* +/-) | Impaired hippocampal inhibitory circuit function | [152] |
| 8q24 deletion;<br>OMIM: 615583 | Craniofacial anomalies, DD, ID, coloboma, cardiac defects, psychomotor delay and convulsions | Contiguous gene syndrome | Zebrafish | Suppression of *SCRIB* and *PUF60* leads to short stature, microcephaly and craniofacial defects. Also, suppression of *SCRIB* leads to coloboma and renal abnormalities | [128] |

ID, intellectual disability; DD, developmental delay; ASD, autism spectrum disorders

## Mouse models

The mouse genome is highly similar to the human genome, with 99% of mouse genes having human orthologs [83]. The mouse genome also displays extensive synteny, with conserved gene order, structure, and orientation compared to the human genome, rendering modeling of rare CNVs in mouse straightforward. Development of a Cre/*loxP* system for targeting chromosomal regions has enabled the creation of different sized deletions and duplications involving disease associated regions [84]. Here, gene targeting is used to insert *loxP* sites into putative CNV breakpoint locations in the ES cell genome, which is then exposed to Cre to induce recombination between the *loxP* sites and generate the rearrangement. These targeted ES cell clones are injected into mouse blastocysts, and embryos are transferred to pseudo-pregnant mice resulting in the birth of chimeras. Chimeric mice with cells targeted for the gene or genomic region of interest are then mated with wild type mice for germline transmission. Mouse models engineered for deletion or duplication of the entire CNV region have been used to recapitulate human CNV disorders, while nested deletions and gene-specific knockouts have been used to map specific genes to a particular phenotype. Genetic mouse models have been generated for a number of rare CNVs, including 7q11.23, 22q11.2, 17p11.2, 15q13.3, and 16p11.2 (Table 1). Some examples are illustrated in this review.

The molecular genetic basis of DGS due to a 22q11.2 deletion, the most common microdeletion disorder, was dissected using mouse models. In patients, this deletion results in craniofacial abnormalities, cleft palate, thymus aplasia (or hypoplasia), hypocalcaemia, and cardiovascular defects [85]. More variable features include cognitive and behavioral defects, such as schizophrenia, intellectual disability, and autism spectrum disorders [86]. In fact, the first causative genes for DGS were identified from mouse studies [87]. Several independent groups identified *Tbx1* as a major contributor to some of the phenotypes observed in DGS patients by using a combination of overlapping deletions and targeted single gene mutations in mice [88–93]. Further analysis of a series of deletions and single gene mutants in mice, using prepulse inhibition as an indicator for sensorimotor gating and behavioral features, confirmed the role of *Tbx1* and implicated the potential role of *Gnb1l* towards psychiatric features associated with 22q11.2 deletion [94]. Only later were mutations in *TBX1* identified in affected human patients confirming their role in DGS phenotypes [94–97]. While these studies have provided insights into the pathobiological basis of DGS, the often-observed high variation in phenotypic expressivity remains unexplained. It is possible that alterations in other genes such as *GNB1l* and *COMT* within or outside of the region [94, 98, 99], along with haploinsufficiency of *TBX1* contribute to variable expressivity. Several genes, both within 22q11.2 and in other genomic regions, have been found to interact with *Tbx1* and exacerbating the phenotypic features in mouse models [99–103]. The results and observations thus far support the model that haploinsufficiency of *TBX1* is sufficient to cause DGS.

Phenotypes observed in SMS and PTLS have also been recapitulated in mice engineered to carry a deletion and duplication, respectively, of the syntenic region on mouse chromosome 11 [104, 105]. The involvement of *RAI1* in SMS was further validated in *Rai1* knockout mice that demonstrated craniofacial and behavioral features [106, 107]. Comparison of SMS mouse models with different sized deletions containing *Rai1* with the *Rai1+/-* mice, in a relatively pure C57BL/6 background, showed that the penetrance of craniofacial features were modified by the genomic region surrounding *Rai1* [108]. Similarly, most features of PTLS were identified in transgenic mice overexpressing *Rai1* [109, 110]. In an elegant study, Walz and colleagues generated compound heterozygous mice carrying both deletion and duplication of 17p11.2 as well as mice carrying the duplication and *Rai1* loss-of-function mutation [111]. The results showed normal Rai1 dosage and rescue of phenotypes due to 17p11.2 duplication, confirming the dosage sensitive role of *RAI1* in both SMS and PTLS.

The molecular genetic basis of WBS has been dissected using mouse models with partial deletions, proximal deletion, distal deletion, double heterozygote and single gene knockouts for genes within 7q11.23 [68, 112-116]. For example, single gene knockouts have been generated for promising candidate genes such as *ELN*, *CYLN2*, *GTF2IRD1*, *GTF2I*, *LIMK1*, *EIF4H*, *BAZ1B*, and *FKBP6* that recapitulated specific phenotypes and mechanisms in WBS. In a recent study, Segura-Puimedon and colleagues generated a mouse model that mimicked the most common deletion found in WBS patients. The complete deletion mouse represented a “true” WBS model, recapitulating almost all of the physical and cognitive defects seen in WBS patients [117]. Mouse models for the reciprocal duplication of WBS on 7q11.23 are not available yet. Mervis and colleagues studied anxiety phenotypes that are observed in about 30% of individuals with the 7q11.2 duplication [118]. Mice that carried increased copies of the general transcription factor 2I (*Gtf2I*) gene produced more separation anxiety behaviors compared to mice with decreased copies of the gene.

Mouse models have also been characterized for certain recently identified rare CNVs. Deletion and duplication of the mouse syntenic region for human 16p11.2 have been created by chromosome engineering [119, 120]. Clinical features in children with the 16p11.2 deletion include autism, intellectual disability, obesity, and epilepsy, while the reciprocal duplication has been associated with schizophrenia and other neurodevelopmental phenotypes. Mice engineered for these CNVs showed behavioral phenotypes and defects in brain architecture with deletion mice showing more severe features than the duplication mice. However, these studies have yet to be extended to identify specific causative genes for the observed phenotypes. Similarly, mouse models of 15q13.3 microdeletion harboring six genes including *CHRNA7* have been created [121]. The 15q13.3 microdeletion has been enriched in individuals with epilepsy, developmental delay, autism, and schizophrenia. Mice carrying deletions of the syntenic region for human 15q13.3 exhibited increased susceptibility to seizures, impaired spatial reference memory, aggression, increased body weight, and auditory processing defects, recapitulating most of the clinical features observed in patients with the deletion. However, certain behavioral phenotypes related to schizophrenia, such as anxiety-related behavior, working memory, and amygdala-dependent learning and memory were not observed in these mice. These phenotypes might be contingent upon the effects of genetic background and/or environmental factors.

## Zebrafish models

Zebrafish is a tropical freshwater fish that has been an attractive model system to study embryogenesis and organ development. The advantages conferred by the short generation time, the ability to evaluate large populations, and easy access of transparent embryos for observation and manipulation make zebrafish a more sought after model for high throughput genomic screens [122, 123]. The annotated genome of zebrafish indicates that about 70% of genes have orthologs in humans [123]. In recent years, the zebrafish has emerged as a powerful model for the functional analysis of genes within rare CNVs. Two landmark studies have exemplified the use of zebrafish for prioritizing candidate genes within 16p11.2. Blaker-Lee and colleagues carefully assessed phenotypes for overexpression and knockdown for all available 16p11.2 orthologs in zebrafish and found that several genes are highly active, contributing to various phenotypes in a dosage-sensitive manner [124]. Using a different strategy, Golzio and colleagues studied epistatic effects by using discrete gene pairs to identify the contribution of specific genes towards neuroanatomical phenotypes [125]. Using morpholinos to suppress the expression of genes and human mRNA injections for overexpression, *KCTD13* was identified as a major driver for the microcephaly and macrocephaly phenotypes associated with 16p11.2 CNVs. Similarly, a recent study on 17p13.1 deletion syndrome showed that dosage alterations of a combination of genes within the critical interval resulted in microcephaly features in zebrafish, reminiscent of those observed in patients. Causative genes responsible for phenotypes in 8q24.3 deletion were also identified using functional studies in zebrafish. Individuals with 8q24.3 deletion manifest craniofacial anomalies, developmental delay, coloboma, cardiac defects, psychomotor delay, and convulsions [126–128]. Using atypical deletions, the minimum critical region was narrowed down to three genes including *SCRIB*, *NRBP2* and *PUF60* [128]. Further functional studies using morpholino-induced knockdown identified the contributions of the cell polarity effector *SCRIB* and the splicing factor *PUF60* towards the phenotypes observed in individuals with 8q24.3 deletion.

## Fruit fly models

*Drosophila* has been used in genetics research for more than 100 years and has been instrumental in understanding gene mutations, genes involved in different pathways, and molecular processes [129]. About 75% of human disease associated genes have orthologs in *Drosophila* [130]. Compared to mouse, the fruit fly carries several advantages, including a short generation time, low costs for rearing and maintenance in the laboratory, and is amenable for behavioral and neuronal studies. There have been several advances in the genetic tool kit of *Drosophila* over the years, including transposition of exogeneous DNA to generate transgenic flies, tissue specific expression using the UAS-GAL4 system [131], and RNA-interference-mediated knockdown [132], among many others. These advances have made *Drosophila* an extremely powerful and sophisticated genetic model system for identifying and analyzing complex biological processes. *Drosophila* also serves as a powerful tool for studying brain development and related disorders as the principles of nervous system development and function are similar to those of vertebrates.

For example, *UBE3A*, the causative gene for Angelman syndrome, has been successfully modeled in fruit flies. Wu and colleagues generated *dUBE3A* null mutants that exhibited motor defects, abnormal circadian rhythms, and long-term memory defects [133]. These results were further reproduced by introducing a transgene containing missense mutations analogous to the ones identified in Angelman syndrome patients in null mutant background. Upon overexpression of the *dUBE3A* specifically in the eye and wing, morphological abnormalities were observed suggesting a significant role in early development. In another study, Lu and colleagues showed that *dUBE3A* null mutants display abnormalities in dendritic arborization of sensory neurons in the peripheral nervous system [134]. They were also able to reproduce these results in RNAi knockdown of *UBE3A*. These findings implicate a role for *dUBE3A* in normal dendritic branching/patterning and suggest that abnormal dendritic arbors may contribute to neurological defects in AS patients. Mutational analysis in *Drosophila* identified a role for *EHMT1* as a regulator of peripheral dendrite development, larval locomotor behavior, non-associative learning, and courtship memory [135]. Interestingly, memory was restored in *EHMT* mutants by restoring the expression of *EHMT* in adulthood, indicating the reversible nature of the cognitive defects associated with *EHMT1* deficiency. Further analysis identified a set of downstream neuronal genes that control learning, memory, and behavior. Thus, fruit flies can be used to dissect specific neuronal mechanisms altered due to mutations in conserved genes within rare CNVs.

## Conclusions

Rare CNVs are like the canaries in the coalmine, pointing toward candidate genes within the regions of interest. Genome architecture predisposes certain regions to recurrent rearrangements, and these regions are significantly enriched for genes important for neurodevelopment. While the causative genes have been identified based on chromosomal events or atypical deletions, individual genes conferring specific phenotypes have not been identified for a vast majority of rare CNVs discovered recently. Even though mice have been the standard models for functional evaluation, the high costs involved in generating and maintaining mouse lines and the daunting task of knocking out each and every gene within the CNV region in mice, among many other reasons, has made way for zebrafish and *Drosophila* as tools for high throughput screening. It is important to note that each model has its own advantages and disadvantages and a combination of all the models is required to understand the molecular mechanisms leading to rare CNV disorders.

## ACKNOWLEDGEMENTS

We thank Andrew Polyak, Claire Reynolds, Lucilla Pizzo, Megan Schilling, and Sneha Yennawar for critical reading of the manuscript.

